# The EFFECT benchmark suite: measuring cancer sensitivity prediction performance - without the bias

**DOI:** 10.1101/2023.10.02.560281

**Authors:** Bence Szalai, Imre Gáspár, Valér Kaszás, László Mérő, Milán Sztilkovics, Kristóf Z. Szalay

**Affiliations:** Turbine Ltd., Budapest, Hungary

## Abstract

Creating computational biology models applicable to industry is much more difficult than it appears. There is a major gap between a model that looks good on paper and a model that performs well in the drug discovery process. We are trying to shrink this gap by introducing the Evaluation Framework For predicting Efficiency of Cancer Treatment (EFFECT) benchmark suite based on the DepMap and GDSC data sets to facilitate the creation of well-applicable machine learning models capable of predicting gene essentiality and/or drug sensitivity on in vitro cancer cell lines.

We show that standard evaluation metrics like Pearson correlation are misleading due to inherent biases in the data. Thus, to assess the performance of models properly, we propose the use of cell line/perturbation exclusive data splits, perturbation-wise evaluation, and the application of our Bias Detector framework, which can identify model predictions not explicable by data bias alone.

Testing the EFFECT suite on a few popular machine learning (ML) models showed that while library-standard non-linear models have measurable performance in splits representing precision medicine and target identification tasks, the actual corrected correlations are rather low, showing that even simple knock-out (KO)/drug sensitivity prediction is a yet unsolved task.

For this reason, we aim our proposed framework to be a unified test and evaluation pipeline for ML models predicting cancer sensitivity data, facilitating unbiased benchmarking to support teams to improve on the state of the art.

## 2 Introduction

Machine learning models are frequently used to predict in vitro drug sensitivity and gene essentiality of cancer cell lines (Adam et al. 2020; Dempster et al. 2020). Interpretation of these models can help to identify sensitivity and resistance biomarkers for different anti-cancer compounds (Dempster et al. 2020) and they promise applicability on in vivo models (Kurilov et al. 2020) and for patient drug response predictions (Jia et al. 2021; Wang et al. 2022). To identify well-performing ML models, correct evaluation metrics and benchmarking are crucial, especially as ML models can learn frequently present data biases (Eid et al. 2021), which makes standard metrics unreliable and confounds model interpretation.

In this study, we focused on predictions made on in vitro data as in vitro cancer cell line sensitivity data is much more abundant than in vivo and patient data. In most cases, more complex in vivo predictions also incorporate in vitro data – with all its corresponding biases. CCLE (Ghandi et al. 2019) and Cell Model Passports (Meer et al. 2019) created extensive multi-omics datasets of >1,000 cancer cell lines, where baseline (unperturbed) gene expression and genomic status of cancer cell lines are available. Other initiatives like GDSC (Iorio et al. 2016), CTRP (Seashore-Ludlow et al. 2015), and PRISM (Corsello et al. 2020) created datasets where drug sensitivity (drug response curve IC50 or area-under-curve (AUC), or their z-score normalized version) is measured in these cancer cell lines for 100s/1,000s of compounds. Complementary, genome-wide gene essentiality measurements are available as the DepMap dataset (Tsherniak et al. 2017). In these experiments, a genome-wide, pooled CRISPR screen was performed in a large panel of cancer cell lines, and gene knock-out-induced viability changes were measured (named gene effect).

What prompted our investigation is the emergence of models simultaneously trained for multiple perturbations (Firoozbakht et al. 2022; Menden et al. 2013; Costello et al. 2014; Yang et al. 2018; Szalai et al. 2019; Manica et al. 2019; Jiang et al. 2022). Typical machine learning models are trained separately for each perturbation (drug or gene knock-out), using the baseline multi-omics parameters of the cell lines as features (Firoozbakht et al. 2022; Iorio et al. 2016; Dempster et al. 2020). In contrast, multi-perturbation methods use the multi-omics parameters of the cell lines together with perturbation-specific features to predict sensitivity for multiple, different perturbations with one model, opening new possibilities like predicting sensitivity for new perturbations unseen during the model training. Despite these rich possibilities, there is still a gap between how high accuracy scores these models can claim, and their lack of adoption in the industry. We argue that this is at least partly due to how far common methods to calculate accuracy are from what happens in actual downstream applications. For example, when the investigation into a new target begins, researchers are interested in the cell lines’ sensitivity to the target of interest before they start synthesizing a compound and they have any training data available for that specific perturbation, so no metric measured on a simple random train/test split will give us the correct picture for this application. We also uncovered additional pitfalls by analyzing the performance of ML models with different input features and complexity on the sensitivity prediction problem. We show that even on an applicable train/test split, standard metrics, like Pearson correlation cannot differentiate between models, as simple models with uninformative features have similar performance to ones using biologically informative features. We identified that cell line and perturbation-specific biases mislead model performance metrics and developed a statistical framework to correct them.

While our framework is informed by our challenges benchmarking cross-perturbation models, we aim for it to be applicable in any type of cancer sensitivity prediction setup. Our goal is to help the field move towards models with better practical use. To further support that goal, we also release our datasets to serve as uniformized benchmark data for further studies at https://benchmark.turbine.ai.

## 3 Results

### 3.1 Prediction and evaluation setup for multimodal machine learning models

To test the benchmarks, we have also built a simple cross-perturbation ML framework, where a single model learned the sensitivity of different cell lines for different perturbations. We have created two separate test schemes: one predicting drug sensitivities, and another predicting gene dependencies. For the current article, we did not mix these two datasets, so separate models were trained for drugs and gene dependencies.

To perform the training, the models require both cell line and perturbation-specific features (**Figure 1A**). We used four different feature setups: a simple cell line identity (one-hot encoding), gene expression, genetic mutations, and the union of all features (Methods) as cell line features. For drug perturbation features, we used drug identity, canonical target, drug target affinity, chemical fingerprint, and again, the union of these features (Methods). In the case of gene perturbations (KOs), we used the gene identity, node2vec-based network topology embedding, prot2vec-based sequence information, Gene Ontology terms of the KO’d gene, and the union of all these features (Methods) as input features.

**Figure 1:**
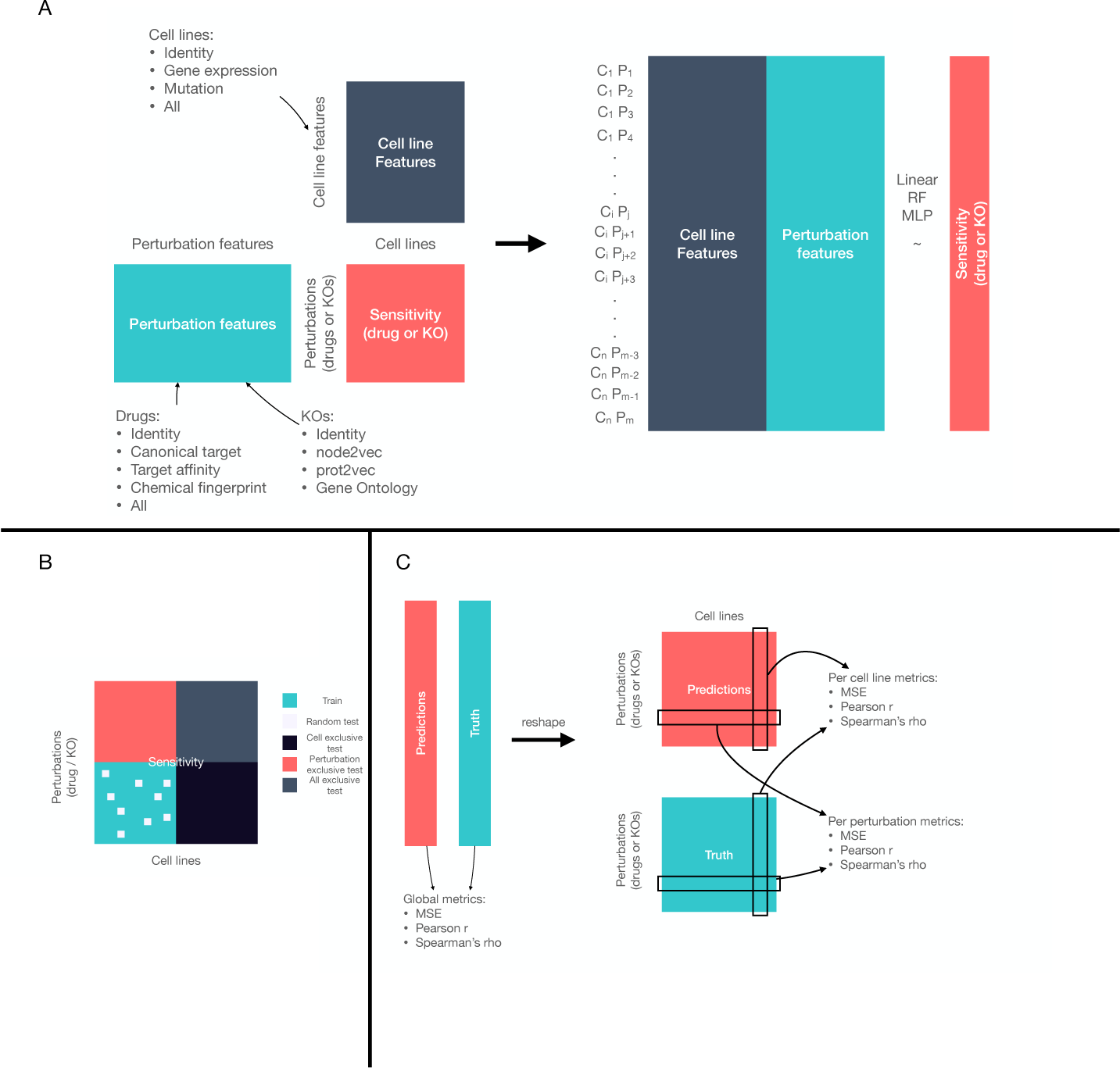
General framework for multimodal machine learning prediction of sensitivity. (A) Model training setup: cell line and perturbation-specific features (left side) are concatenated for each sample (unique cell line – perturbation pair) and are used to predict target sensitivity variable (gene effect or drug IC50). (B) Composition of training and test sets. (C) Evaluation metrics: global metrics are calculated for the whole predicted dataset (left) while per-cell line/per-perturbation metrics are calculated for each of the respective subsets of predictions.

For each sample (cell line – perturbation pair), the corresponding cell line and perturbation features were concatenated, and the final feature matrix was used to predict drug sensitivity (IC50 or z-score) from GDSC (Iorio et al. 2016) or gene essentiality (gene effect) from DepMap (Meyers et al. 2017; Behan et al. 2019; Dempster et al. 2019), using ML models with different complexity; linear regression (LR), Random Forest regression (RF), or Multilayer Perceptron regression (MLP). We used separate models for drug sensitivity and gene essentiality predictions.

As highlighted previously, one of the most important considerations when setting up a model is to select the right train/test splitting regimen for the downstream application. Just holding out a random set of genes (or drugs) and cell lines from the train set already yields three different test sets, each much more representative of a given downstream application than a simple random split (**Figure 1B**). The CEX (cell-exclusive test) slice maps to an application where one wants to predict the performance of a known drug in new models, such as in drug repurposing or precision medicine.

The DEX or GEX (drug or gene-exclusive, depending on the type of perturbation, together also referred to as perturbation-exclusive, PEX) slice maps to target discovery applications – when we would like to assess the effect of a new drug on a set of established in vitro model cell lines. For testing purposes, we did include a random train/test split as well (RND). Notably, we could not think of practical downstream applications for the easiest (RND) and the hardest (AEX - all exclusive) splits, so we would advise against using these to generate the primary metrics in benchmarking scenarios. To monitor the variability of prediction performance, we created 3 sets of random train-RND-CEX-PEX-AEX splits (Methods).

To show which metrics are the most informative, we used common metrics for the evaluation of model performance (**Figure 1C**). Mean-squared error, Pearson, and Spearman’s correlation between the full prediction and truth vectors were used as global metrics. We also calculated these metrics for each cell line and each perturbation separately and called these metrics cell line and perturbation-wise metrics.

The different ML model types, featurization, and test sets allowed us to define control models that were expected to perform badly before the experiments. For example, as modeling different perturbations in cell lines requires learning interactions between cell line and perturbation features, we expected that linear models would not perform well in these setups and were used as control experiments. Similarly, identity (one-hot encoding) features are not expected to perform well in some of the test setups, namely identity cell line features in CEX and AEX setup and identity perturbation features in PEX and AEX setup, so these models are also control experiments.

This experiment setup enabled us to get information about a wide range of model types, train/test splits, and evaluation metrics, enabling us to analyze model performance in an unbiased way.

### 3.2 Biases of sensitivity data

Machine learning models that use high-dimensional data can easily learn trivial biases/confounding factors of datasets. Learning these biases can increase apparent model performance, however, it can hinder model generalization and lead to false interpretation and biological translation of results. We identified that most of the biases are related to the general sensitivity of cell lines to perturbations and the general effectiveness of perturbations and developed a statistical framework to overcome this problem.

Different drugs have different mean (median) IC50 values (**Figure 2A**). This difference is mainly attributable to the chemical properties and target profile of drug molecules. While mean IC50 is important from a chemical feasibility point of view, it is less important and even can be misleading when assessing biological selectivity. For example, the cell line CORL88 has a lower IC50 for Bortezomib than for Cyclophosphamide (**Figure 2A**), but its Cyclophosphamide IC50 is in the lower range of the whole Cyclophosphamide IC50 distribution, making its relative sensitivity higher for Cyclophosphamide than for Bortezomib. This drug bias can also confound the evaluation of prediction models: a model only learning the mean IC50 of drugs (with some random noise), can already show strong prediction performance using standard correlation metrics (**Figure 2B**, Pearson correlation *r* = 0.96).

**Figure 2:**
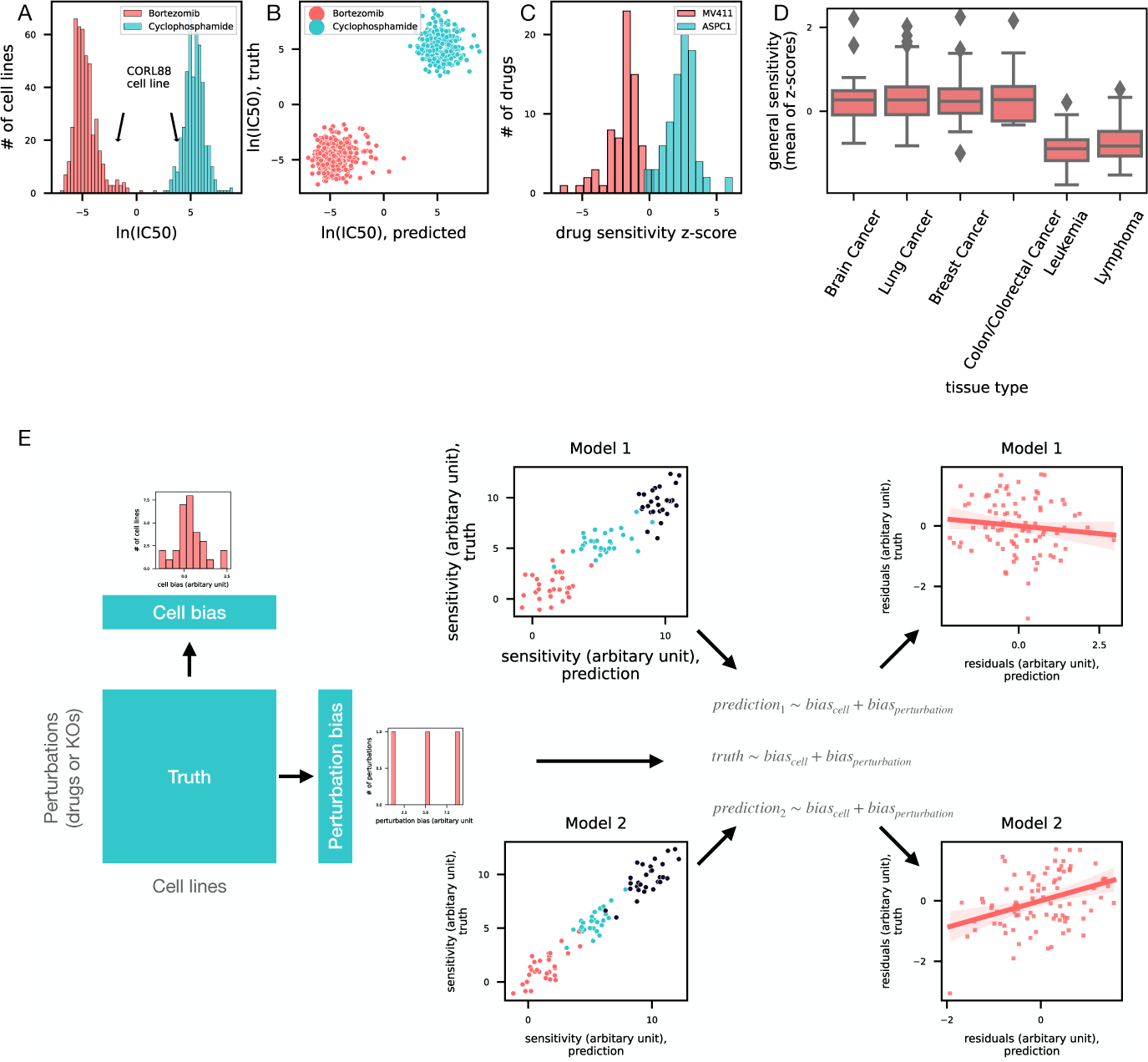
Biases of sensitivity data and Bias Detector (A) Mean logIC50 as confounding factor (B) Effect of mean logIC50 bias on evaluation metric (C) General sensitivity of cell lines as confounding factor (D) Tissue type distribution of general sensitivity (only 6 most frequent tissue types are shown) (E) Bias detection framework: Two models (Model 1, which learned only biases with some error, and Model2, which learned some additional information) have similar performance in predicting sensitivity based on the raw Pearson correlation as a metric (middle, Pearson correlations 0.92 and 0.96, respectively, color code represents 3 different drugs of this toy example). Cell line and perturbation-specific biases are calculated from truth data (left). A linear model is fitted for truth and predicted values, and Pearson correlation is calculated between residuals (partial correlation coefficient, right). Bias detector identifies the increased performance of Model2 (partial correlation coefficients and p values: −0.11, 0.29, and 0.37, 0.0002 for Model1 and Model2, respectively).

To overcome this problem, relative sensitivity (z-score) is a frequently used target metric in drug sensitivity prediction. Z-score is calculated by drug-wise normalization of log(IC50) values, thus removing the confounding effect of mean drug IC50. However, cell lines also have intrinsic biases, like general sensitivity (Geeleher et al. 2016), meaning that some cell lines are sensitive (MV411 cell line), while others are resistant (ASPC1 cell line) to all investigated drugs countering the effect of z-score normalization (**Figure 2C**). General sensitivity can be a consequence of drug efflux transporters but can also arise from different division times of cell lines (Hafner et al. 2016). Importantly, general sensitivity is associated with tissue type (**Figure 2D**), thus it can be learned by machine learning models (e.g., from gene expression features) and can confound model evaluation just like the previously mentioned mean drug IC50 bias. Similarly, perturbation biases can also be learned from biologically inspired features.

While using per-cell line and per-perturbation metrics (**Figure 1C**) can partially overcome these problems, these evaluations remove only one of these biases at once (per-cell line evaluation removes cell bias, and per-perturbation evaluation removes perturbation bias). To overcome this problem, we created a statistical framework, named Bias Detector (BD) to systematically remove both cell line and perturbation-specific confounding factors, and evaluate ML model performance in a more unbiased way. We calculated cell line and perturbation biases for the whole GDSC or DepMap dataset (Methods). Then, we calculated partial correlation coefficients between predicted and true values, using the pre-calculated cell line and perturbation biases as covariates (**Figure 2E**). Our final reported metrics are the partial correlation coefficients and the fraction of perturbations (or cell lines) where the partial correlation of the predicted and true values is significant. We argue that this BD framework can identify models that learn specific and biologically interesting sensitivity patterns, not only trivial cell line or perturbation biases.

### 3.3 Predicting gene essentialities

We first subjected models trained to predict gene essentiality to our proposed evaluation framework. This showed that using global correlation metrics, even the simplest model (linear regression trained using one-hot encoded feature set) has an astoundingly good performance (Pearson *r* ≈ 0.8) not only in the RND split but also in the cell-exclusive (CEX) split, despite one-hot encoded cell features not being expected to convey any generalizable message (**Figure 3A and 3B**). This is in strong contrast to correlation scores calculated per gene, where few genes score above 0.3 in any setup except the MLP model on the random split (**Figure 3D**) in line with prior expectations. Furthermore, in the CEX split, all linear regression models, and even some of the more complex setups, lost most of their predictive performance. Only models trained on the most information-rich feature sets (**Figure 3E**) were predictive in this harder, but more practical generalization scenario. These observations agree with the hypothesis that good global correlations could be reached by only learning the average sensitivity of cell lines to different perturbations and nothing else.

**Figure 3:**
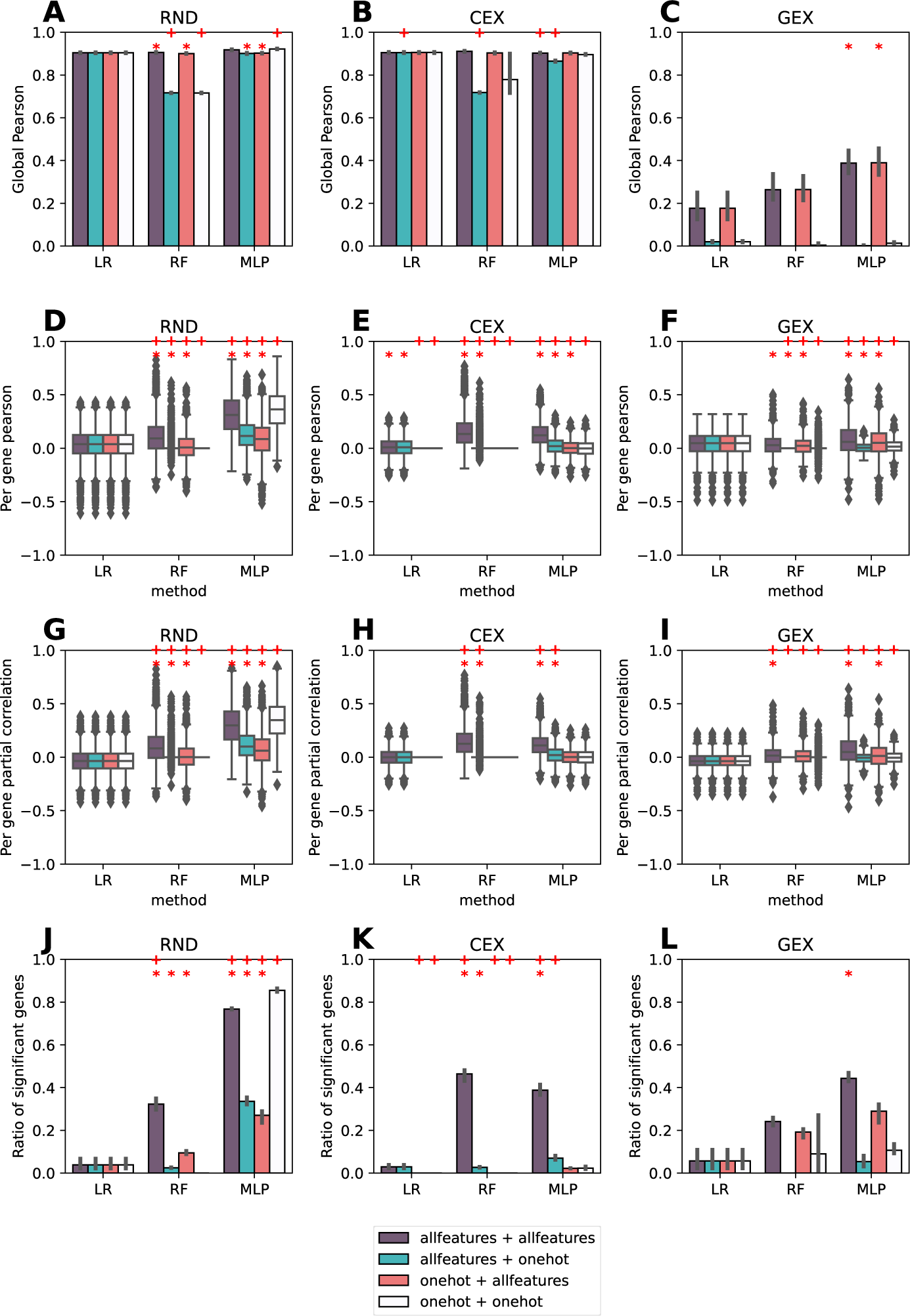
Model performance of predicting DepMap gene effect. (A-C) Raw Pearson correlation of model predictions to ground truth pooled from three different RND (A), CEX (B), and GEX (C) splits. (D-F) distribution of Pearson correlations calculated separately for each perturbed gene for the three splits: RND (D), CEX (E), and GEX (F). (G-I) Distribution of per perturbation partial correlations as reported by BD for the three different splits. RND (G), CEX (H), DEX (I). (J-L) Fraction of perturbations with significant positive partial correlations as detected by BD; mean and 95% confidence interval of three different RND (J), CEX (K), and DEX (L) splits. Different colors indicate different cell line + perturbation feature set combinations (legend). The first part signifies cell features, the second perturbation features. Red plus sign indicates significant difference (*p* <= 0.05) compared to LR trained on all cell and perturbation features (meaning improving the algorithm was significant), red asterisk indicates significant difference (*p* <= 0.05) compared to the respective method trained on one-hot encoded cell and perturbation features (meaning improving the data was significant) (see Methods).

The gene-exclusive (GEX) split fared significantly worse in terms of global correlation (**Figures 3C and 3F**). Here, even global correlation can somewhat differentiate between feature sets suitable or unsuitable for generalization. We also saw (like with the other two split types) that the per-gene correlations are greatly reduced compared to the global values. This suggests that models can predict the general essentiality of unseen genes, but they are unable to give information about which cell lines will be more sensitive to a given perturbation. The per-cell line correlations for the GEX set are also high, further proving this hypothesis (Supplementary Figure 1).

Applying the Bias Detector (BD) workflow (partial correlation between model predictions and ground truth, see **Figure 2E** and Methods) to assess the effect of learning general sensitivity on the model performance showed that those models that demonstrated some positive predictive power per gene (i.e., beyond predicting the mean difference between the effect of individual genes) were predicting more than general sensitivity for some genes (**Figures 3G to 3L**). Importantly, this was also only possible if information-rich feature sets were used for model training. Without these measures, the user of a system wouldn’t have been able to understand which genes’ predictions are reliable enough to base their in vitro experiments on.

### 3.4 Predicting drug sensitivity

In contrast to predicting gene effect, all three models predict drug sensitivity – both IC50 (**Figure 4A**) and z-score (Supplementary Figure 2) – well in the RND split as indicated by the high Pearson correlation coefficients per perturbation. Interestingly, linear regression gave a very consistent performance, irrespective of the cell and perturbation features used (**Figure 4A** and Supplementary Figures 3 and 4). As expected, none of these three methods could predict drug sensitivity in the CEX split when trained only on the OHE (one-hot encoded) cell features (**Figure 4B** and Supplementary Figure 3). Surprisingly, however, all methods achieve high per-perturbation scores in the DEX splits, even when trained on OHE perturbation features, which are expected to be non-generalizable (**Figure 4C** and Supplementary Figure 2). This suggests that the models likely learned some trivial feature of the dataset, e.g., the general sensitivity of cell lines – which in the DEX split were all part of the training set also – to any perturbations. This cross-contamination of the measure by bias factors orthogonal to the test feature is exactly why we needed a separate Bias Detector.

**Figure 4:**
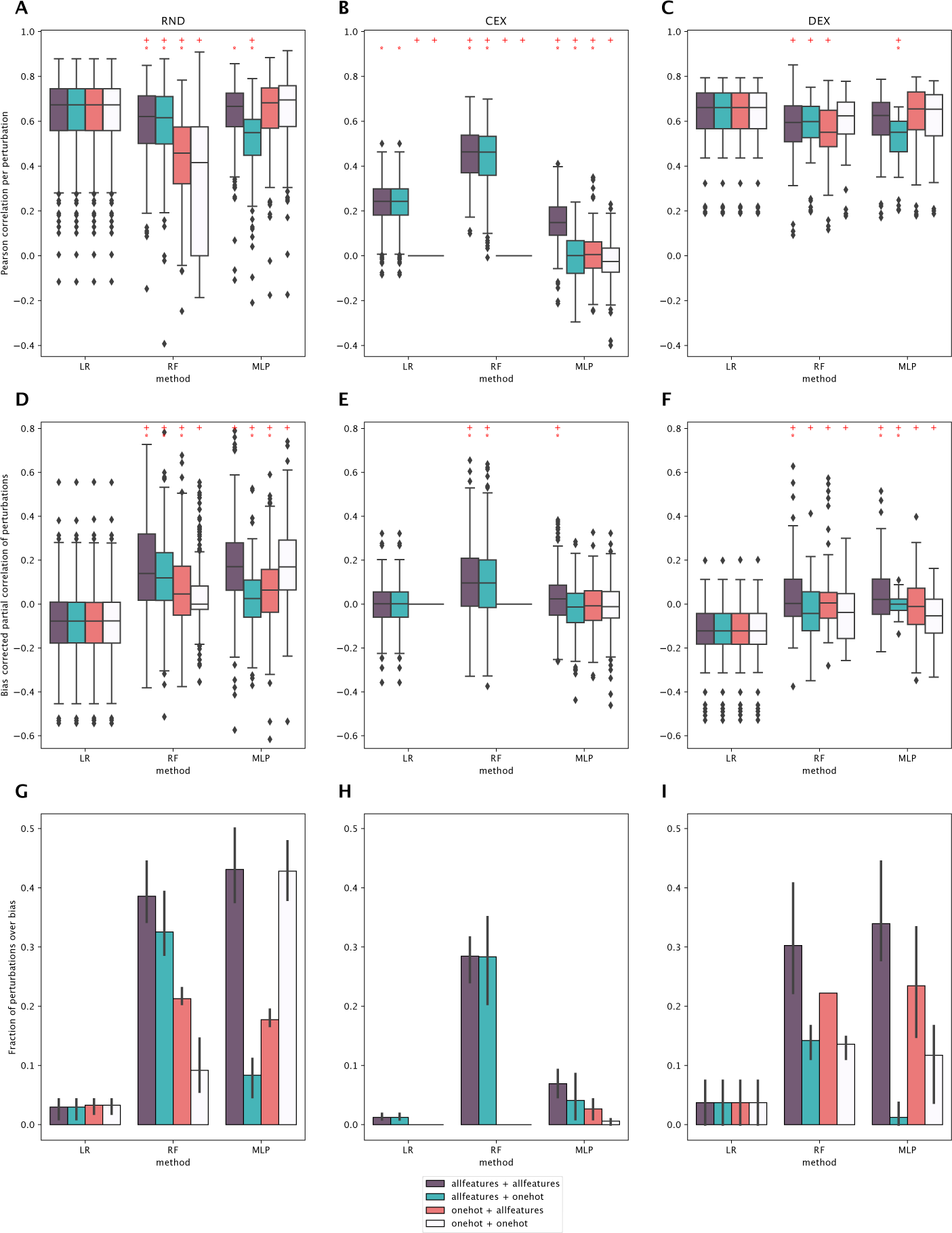
Model performance of predicting IC50 drug sensitivity. (A-C) distribution of Pearson correlations calculated separately for each perturbed gene for the three splits: RND (A), CEX (B), and DEX (C). (D-F) Distribution of per perturbation partial correlations as reported by BD for the three different splits. RND (D), CEX (E), DEX (F). (G-I) Fraction of perturbations with significant positive partial correlations as detected by BD; mean and 95% confidence interval of three different RND (G), CEX (H) and DEX (I) splits. Different colors indicate different cell line + perturbation feature set combinations (legend). The first abbreviation signifies cell features, the second perturbation features. Red plus sign indicates significant difference (*p* <= 0.05) compared to LR trained on all cell and perturbation features (meaning improving the algorithm was significant), red asterisk indicates significant difference (*p* <= 0.05) compared to the respective method trained on one-hot encoded cell and perturbation features (meaning improving the data was significant) (see Methods).

To evaluate whether such confounding factors could explain the unexpectedly good performance, we used the Bias Detector to calculate partial correlation coefficients and the fraction of perturbations that are predicted with significantly good partial correlation to ground truth (adjusted *p* < 0.05, see **Figure 2E** and Methods) just like before. This showed that linear models largely just learned the bias that exists in the drug sensitivity dataset as very few perturbations were found to be significantly better than the bias correlation by the BD **Figures 4D to 4I**). This also explains the consistent performance of the linear regression models irrespective of the feature sets used for training (**Figures 4A and 4C**). Non-linear models, random forest in particular, were able to provide predictions beyond the data bias; in the RND split, the number of perturbations predicted better than data bias largely co-varies with the raw model performance (**Figures 4A and 4G**). This co-variance can be observed to some extent also for the CEX and DEX splits. Most importantly, in the DEX split, we detected a substantial drop in the fraction of over-bias predicted perturbations when OHE perturbations were used to train the non-linear models (especially prominent when predicting z-scores, **Figure 4I** and Supplementary Figure 2). This indicates that even the non-linear models learned mainly trivial associations of drug sensitivity without proper perturbation features. Taken together, evaluating our models trained to predict drug sensitivity revealed that, similarly to the models trained to predict gene effect, a) linear models may have promising raw performance metrics, yet have limited capacity to learn beyond the bias existing in the data and b) non-linear models, even when having poorer raw metrics than LR, can learn non-trivial associations when trained on generalizable features.

Based on the substantial differences between the raw and Bias Detector-based prediction performances, we were interested in which drugs can be better predicted using raw metrics (per-perturbation Pearson correlation) and BD metrics (partial correlation). We performed pathway enrichment using the drug target–prediction performance (correlation or partial correlation) vectors. Our results show (Supplementary Table 1) that general cytotoxic drugs (targeting pathways like Cell cycle, G1 Phase, and caspase activation) are dominating the top results for raw correlations while targeted therapies (targeting pathways including RAF activation and ERBB signaling) are showing the best partial correlations. These results suggest that for general cytotoxic drugs, predicting drug-specific effects is harder, while in the case of targeted therapies, lower raw correlations can be achieved, also underlying the importance of the BD framework.

## 4 Discussion

In this study, we presented considerations to create more effective benchmarks. We showed that data biases can significantly influence model performance and may render the distinction between models using uninformative and informative features difficult. We presented a statistical framework named Bias Detector, which accounts for the effects of cell line and perturbation biases on model performance and can help find the best-performing models.

We showed the importance of choosing the right train/test split to use for benchmarking to make sure the results translate to the downstream applications. In the case of random splits, every model trained on any feature combination reached a comparable, good global correlation with ground truth data, which would suggest that the prediction of in vitro cancer sensitivity is a simple task. Yet, when analyzing the model performance on individual perturbations, model performance dropped quite substantially, in certain cases – e.g., linear regression on predicting the effect of CRISPR KO perturbations – close to a mean of zero correlation. This indicates that ML models trained on a large corpus of perturbations may indeed only learn the mean differences between the effect of those perturbations while failing to sort the samples per perturbation correctly. Linear regression is a prime example of such a bad-performing model class in predicting the CRISPR KO effect; interestingly, linear models trained on any feature combination performed well in predicting drug perturbation effects – even superior to their non-linear counterparts. However, when subjected to the Bias Detector, linear models turned out to be largely incapable of providing better predictions than what the data bias alone allows for – which also explains the feature insensitivity of linear models. Taken together, these observations on the commonly accepted random train/test splits substantiate the better evaluation of ML models trained on large corpora of heavily correlated data such as cancer sensitivity.

Cell exclusive splits represent a therapeutically more relevant scenario, where the best performing existing drug (or drug target) needs to be selected for a given sample representing a patient group (with the strong caveat that actual patient cells are quite unlike any in vitro cancer cell line). Again, using global evaluation metrics, models had similar performance. In a CEX setup, the general effectiveness of a perturbation can be easily learned from the train set, and as general perturbation effectiveness dominates the sensitivity data (**Figure 2A**), this model- and feature-independent performance is not surprising. In the case of perturbation-wise evaluation, we saw an increased performance of the Random Forest model trained on cell features containing biological information (gene expression, in particular), compared to linear and MLP models. Interestingly, mutation features were not as effective cell line features as transcriptomics, which agrees with the results of previous studies (Dempster et al. 2020; Iorio et al. 2016). Using perturbation-wise evaluation, the MLP model had a similar, or even weaker performance than the baseline linear model. Nevertheless, the Bias Detector did reveal that MLP was able to predict better than data biases for more perturbations (and cell lines) than linear models, also highlighting the importance of bias-independent evaluation metrics. Based on our testing, the CEX scenario, while being a stricter test than the pure random split, is still the easiest of the exclusive splits.

Gene/drug exclusive splits represent another application, target identification, that is, predicting sensitivities for a new drug candidate on existing cells. In our opinion, this setup comes closest to the real-life application of an ML model predicting in vitro cancer cell sensitivities. In this test, we already saw differences even in the global metrics: models using biologically informative perturbation features had in general better performance than the ones with one-hot encoded features. In the DEX drug sensitivity predictions, the per perturbation scores were also reasonably good (even in the case of a linear model and uninformative features) suggesting an orthogonal, confounding bias creeping in. We assumed that this confounding bias was the general level of drug sensitivity ((Geeleher et al. 2016), **Figures 2C and 2D**), which was already shown to have a major contribution to apparent model performance. This observation was what originally prompted the development of the Bias Detector tool, which indeed successfully revealed that only non-linear models, using informative features can learn additional information. These effects were less pronounced in the case of gene essentiality prediction, as the normalization methods used by DepMap mostly remove these cell line-specific general effects (Dempster et al. 2021).

As an aside, analyzing the best-performing features showed that GO encoding was the best-performing feature for gene effect, and drug target encoding for drug sensitivity prediction.

For completeness, we also performed experiments in an all-exclusive (AEX) setup, where both the cell lines and the perturbations were new (out of distribution) for the models. In the case of global metrics, we observed similar behavior to the GEX/DEX setup, regarding per-node and per-cell metrics, and after Bias Detector, none of the models showed predictive performance (Supplementary Figure 1). This exemplifies quite well how hard actual clinical drug discovery is when treating new patients with new drugs^1^.

While this first version of the EFFECT suite only uses continuous metrics, it should be noted that the same sensitivity to bias applies to standard classification metrics like accuracy, F1 score, and AUROC. The same tweaks we recommend to correlations could also be applied to these other scores to yield more translatable results (mean per-node scoring, performance over bias).

A natural extension of these cross-perturbation ML models is predicting sensitivity for combinatorial perturbations (Menden et al. 2019), including the application of whole cell models (Yuan et al. 2021; Nilsson et al. 2022). The biases identified in our study are possibly also present in these scenarios, and the applied benchmarking and bias removal tools can be potentially extended for them.

Combinatorial prediction setups naturally have more sources of bias, which highlights something that is fairly trivial, but still important to mention: while any known bias terms can be added to the Bias Detector’s regression logic so that it can account for those, it does not protect the user against unknown biases which are not in the expressed logic. In fact, what is meaningful data and what is bias depends on the actual application. For example, whether the indication (the tissue type of the originating cell line) is a bias depends on the phase of drug discovery the model is being used in. Early on, when trying to find a target indication for a drug of interest, the tissue type is core information. Later, if one is already using a model to recruit for a clinical trial with a pre-set indication, the same tissue type becomes a bias to control for.

We believe that our Bias Detector framework identifies the most important data biases of sensitivity prediction in a scenario where it would be used downstream for finding target indications, but it can be extended to additional confounding factors/data biases.

In summary, we created an extensive benchmark for gene effect and drug sensitivity prediction ML models. We showed the importance of different test splits and evaluation metrics and developed the Bias Detector, a statistical framework to identify data bias-independent, biologically interesting predictive models. Our general recommendation is therefore threefold: 1.) choose the train/test split wisely for the application, 2.) use local, perturbation- or cell-wise metrics instead of global metrics, and 3.) use statistical tools, like our bias detector Bias Detector reduce the effects of data biases in the performance evaluation.

To make these considerations easy to apply, we plan to make the presented benchmark datasets and methods available to the research community at https://benchmark.turbine.ai and maintain and extend them in the future hoping that they can be used for a more unified evaluation of ML models in the field. The very next version we plan to release is one where the drug and gene test sets are harmonized so that models trained on both drug and gene dependency data can be evaluated as well.

## 5 Methods

### 5.1 Dataset composition

We used GDSC2 (ln(IC50) and z-score) data (Iorio et al. 2016) as the base drug sensitivity, and DepMap CRISPR data (Meyers et al. 2017) as the base gene essentiality dataset. In the case of the gene essentiality dataset, we have downsampled the dataset since the large number of non-essential genes would hinder model fits. We labeled each cell line – gene KO pair as DEAD if the gene effect score was below –0.5. We assigned genes to 20 evenly spaced bins according to the ratio of DEAD cell lines for the given gene’s KO. After this step, we randomly selected 200 genes at most from each bin. Bins with less than 200 genes were fully included in the dataset. For the DepMap CRISPR data, 20% of available genes and 20% of available cell lines were split off to form the exclusive test sets. The test cell lines combined with train genes were used for the GEX set, while test cell lines combined with train genes were used for the CEX set. The RND split was taken from the rest of the data points. To ensure the reproducibility of results the random sampling of test genes, cell lines, and samples was repeated three times.

### 5.2 Generating input features

#### 5.2.1 Cell line features

We used gene expression (log-transformed TPM values) and mutation (binary indicator matrix) data from DepMap as descriptive cell line features. To reduce dimensionality, Principal Components Analysis (PCA) was performed, and the first 950 and 671 components, respectively, explaining 90% of the total variance were kept as features.

#### 5.2.2 Drug features

The following features were assembled to describe drug features for the model training: a) One-hot encoding of drug identity, b) Target encoding, where the GDSC annotated target gene(s) or structures (etc. microtubules, chromatin) were encoded in a ternary indicator matrix (1 for inhibitory, −1 for activator role, 0 – for no effect on target), c) Target affinity, similar to target encoding but the matrix values are the log-transformed affinity values of the compound to its targets as reported in BindingDB (Gilson et al. 2016), and d) PubChem fingerprints based on SMILES in a one-hot encoded format, reduced by PCA from 1k to 75 features retaining 90% of the variance. Based on feature availability, 141 drugs from GDSC2 were kept for training and test purposes.

#### 5.2.3 KO-specific perturbation features

We used the following features to represent the target (KO’d) node in CRISPR perturbation experiments: a) One-hot encoding of the node identity. b) Molecular function and biological process Gene Ontology (The Gene Ontology Consortium 2019) terms of the target node (GO features were one-hot encoded and dimension reduced by PCA). c) node2vec encoding of the target node. We used the OmniPath (Türei et al. 2021; Türei et al. 2016) network and the node2vec algorithm (Grover and Leskovec 2016) to create signaling network-based embedding of the target node. And d) Protein sequence embeddings from UniProt (Coudert et al. 2023).

### 5.3 Machine learning model training

The training inputs were given as concatenated feature vectors of various lengths depending on the number of included feature types. We applied three types of models to these datasets: ridge regression, random forest regression, and multi-layer perceptron.

For the implementation of the ridge regression models, we used the Scikit-learn software package (Pedregosa et al. 2011). Ridge regression is a linear model that is regularized through the L2-norm of the model parameters. We chose ridge regression as it is a widely used model for supervised prediction of omics datasets and from a theoretical perspective it should handle well the large number of multicollinear features of our datasets (Wieringen 2015). We optimized the model through the minimization of the mean-squared error (MSE) loss between the cell sensitivity labels and the model predictions. We used the train split of the dataset to do a hyperparameter selection process for the strength of the regularization parameter based on cross-validation. Finally, we applied the selected model for the required test splits.

As another benchmark algorithm, we used the “randomforest” model from the Scikit-learn software package. Random forest is a frequently used model type for life science applications, and we used it as a black box solution without the need for parameter optimization (Touw et al. 2013). We optimized the model through the minimization of the mean-squared error (MSE) loss between the cell sensitivity labels and the model predictions.

In our experiments, a standard multi-layer perceptron (MLP) with rectified-linear unit (ReLU) activation was chosen as a strong general baseline architecture, to model the underlying complex, non-linear relationship between perturbations and cell line responses. After the processing of multiple fully connected layers, a final output layer consisting of only a single neuron with linear activation returns the prediction value for the regression task. The optimization criterion was to MSE loss between the ground truth label and prediction. To avoid overfitting, dropout was applied on the hidden layer neurons, and the best-performing model was selected from the epoch of lowest MSE loss on the random split of each feature collection setup. The optimization of the network parameters was done via back-propagation, using Adam optimizer with *learning rate* = 10^−3^ and *batch size* = 256, *epochs* = 20. The optimal architecture with *depth* = 3, *hidden neuron count per layer* = 128 and *dropout rate* = 0.2 was selected after an extensive hyperparameter search. The source code implementation is provided in the GitHub repository: https://github.com/turbine-ai/DrugKOPred-benchmark-bias.

### 5.4 Pathway enrichment

We calculated Reactome (Jassal et al. 2020) pathway enrichment on the target–correlation coefficient vectors (either raw or partial correlations, perturbation-wise calculation) using the viper (Alvarez et al. 2016) function of decoupleR (Badia-i-Mompel et al. 2022). Correlation values were at first aggregated target-wise.

### 5.5 Model performance evaluation, Bias Detector, and statistical analysis

For each prediction experiment (predications on a specific test set), we calculated global, perturbation, and cell line-wise metrics. For global metrics, we calculated root-mean-squared error (RMSE), Pearson correlation, and Spearman’s rank correlation between the whole prediction and truth vectors (all cell lines and perturbations). For perturbation-wise and cell-wise evaluation, we calculated the same metrics for each perturbation/cell line separately.

For the Bias Detector framework, we calculated cell line and perturbation bias as a first step. For bias calculation, we fitted a *sensitivity_c,p_ ∼ cell_c_* + *perturbation_p_* linear model, where cell and perturbation factors represent the cell lines and perturbations, respectively. We used the coefficients of this linear model as biases. To evaluate the prediction performance of a model, we fitted two linear models: *y pred_c,p_ ∼ cell bias_c_* + *perturbation bias_p_* and *y true_cp_ ∼ cell bias_c_* + *perturbation bias_p_*. The Bias Detector calculated the Pearson correlation and p-value between the residuals of these two models.

The significance of differences in model performance metrics was evaluated statistically by non-parametric Kruskal — Wallis (KW) tests followed by pairwise Mann—Whitney U tests as the metrics themselves are not expected to be normally distributed due to their limited range (−1 to 1 for correlation metrics, 0-1 for fraction of perturbations/cells that are predicted better than the bias model). The testing was done on a per split basis, first evaluating if there was a significant difference (*alpha* = 0.05) in the metrics of all the models trained on any of the analyzed feature combinations by KW test followed by comparing all models by pairwise U tests to the corresponding linear regression model trained on the full combination of descriptive features (ALL+ALL feature set). Furthermore, the effect of the feature set on the model metrics was evaluated similarly, on a per model per split basis: KW test was used to determine the significant difference among the same type models trained on different feature sets and if this test was significant (*alpha* = 0.05), the models were compared by pairwise U tests to the corresponding model trained on the one-hot encoded feature set. P-values of the U tests were adjusted for multiple testing by the Bonferroni-Hochberg method and significance was reported at (corrected) *alpha* = 0.05 level.

## 6 Acknowledgements

We thank Andreas Bender, Aviad Tsherniak, and Daniel V. Veres for the helpful discussions regarding the manuscript.

## 7 Code and data availability

All data and code to reproduce our analysis and to use the evaluation framework is available at https://benchmark.turbine.ai and https://github.com/turbine-ai/DrugKOPred-benchmark-bias, respectively.

## 8 Author contribution

Bence Szalai: Conceptualization, Analysis, Writing, Project administration. Imre Gáspár: Methodology, Analysis, Writing. Valér Kaszás: Software, Writing. László Mérő: Methodology, Writing. Milán Sztilkovits: Software, Writing. Kristof Z. Szalay: Conceptualization, Resources, Writing, Supervision.

## Supplementary information

**Supplementary Figure 1:**
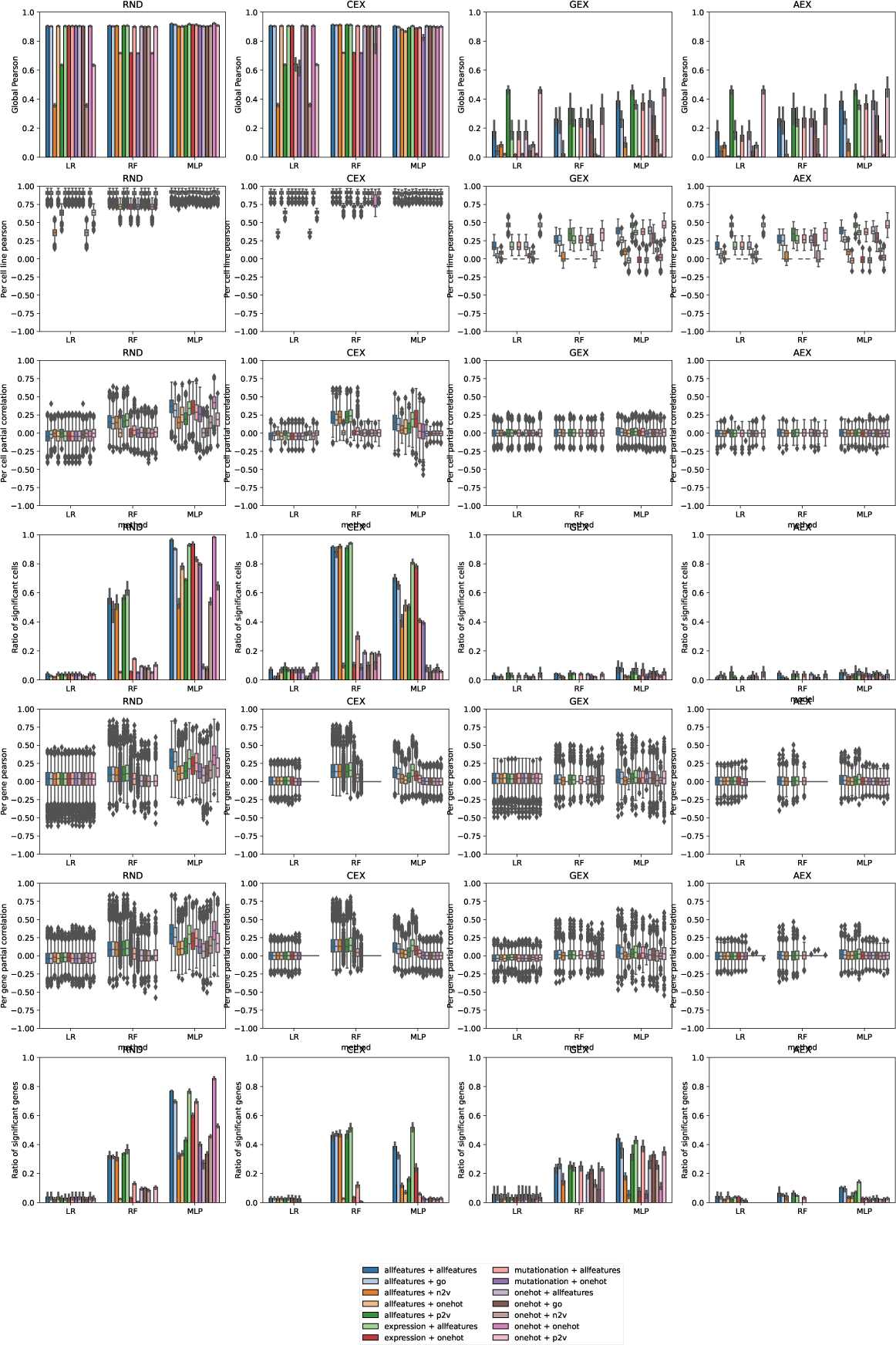
Model performance of predicting DepMap gene effect. Used evaluation metrics are labeled on the y-axis, used ML model is labeled on the x-axis, results are grouped based on splits (RND, CEX, GEX, and AEX columns). Used cell line and perturbation feature pairs are color-coded (legend).

**Supplementary Figure 2:**
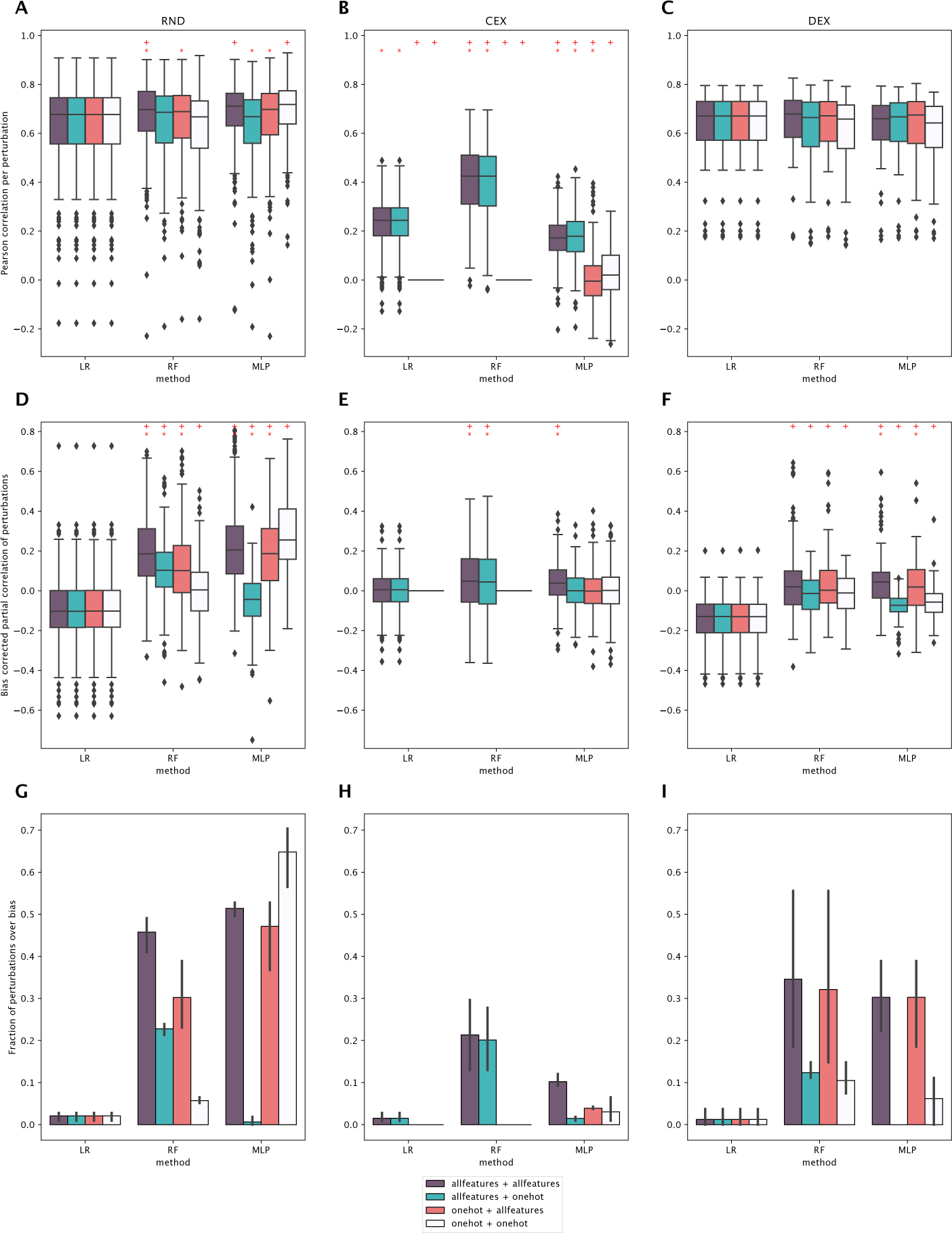
Model performance of predicting z-score drug sensitivity. (A-C) distribution of Pearson correlations calculated separately for each perturbed gene for the three splits: RND (A), CEX (B), and GEX (C). (D-F) Distribution of per perturbation partial correlations as reported by BD for the three different splits. RND (D), CEX (E), DEX (F). (G-I) Fraction of perturbations with significant positive partial correlations as detected by BD; mean and 95% confidence interval of three different RND (G), CEX (H) and DEX (I) splits. Different colors indicate different cell line + perturbation feature set combinations (legend). The first abbreviation signifies cell features, the second perturbation features. Red plus sign indicates significant difference (*p* <= 0.05) compared to LR trained on all cell and perturbation features (meaning improving the algorithm was significant), red asterisk indicates significant difference (*p* <= 0.05) compared to the respective method trained on one-hot encoded cell and perturbation features (meaning improving the data was significant) (see Methods).

**Supplementary Figure 3:**
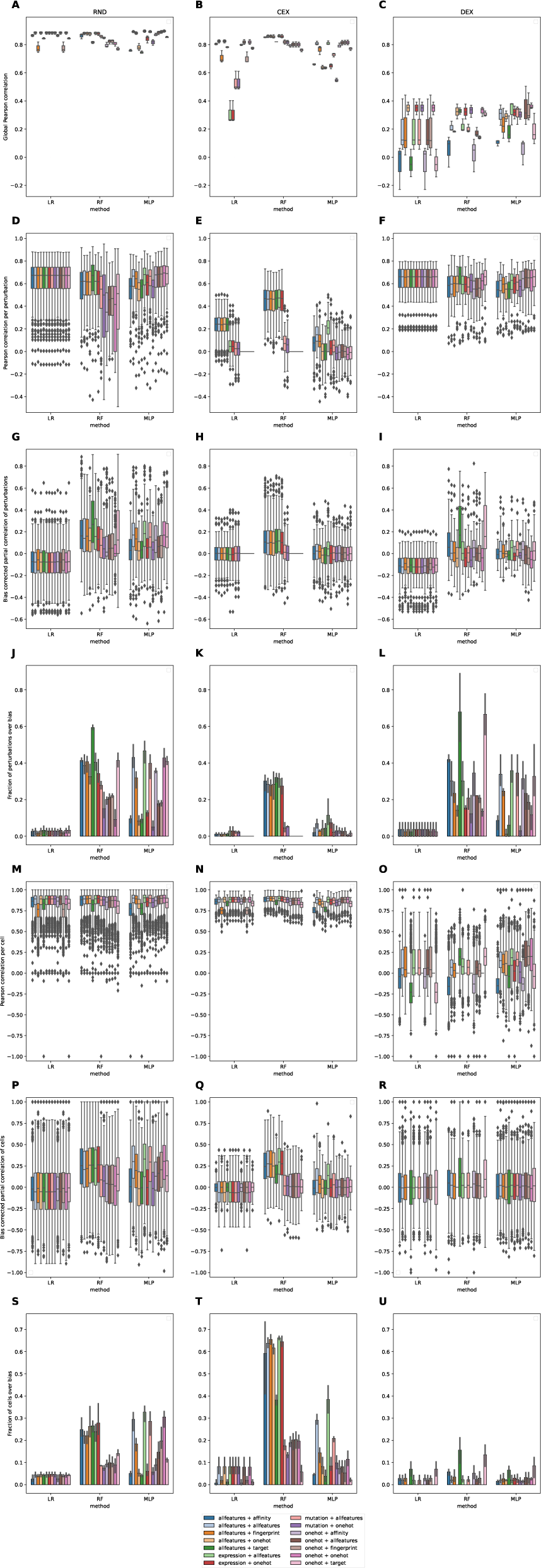
Model performance of predicting IC50 drug sensitivity. Used evaluation metrics are labeled on the y-axis, used ML model is labeled on the x-axis, results are grouped based on splits (RND, CEX, GEX, and AEX, columns). Used cell line and perturbation features pairs are color-coded (legend).

**Supplementary Figure 4:**
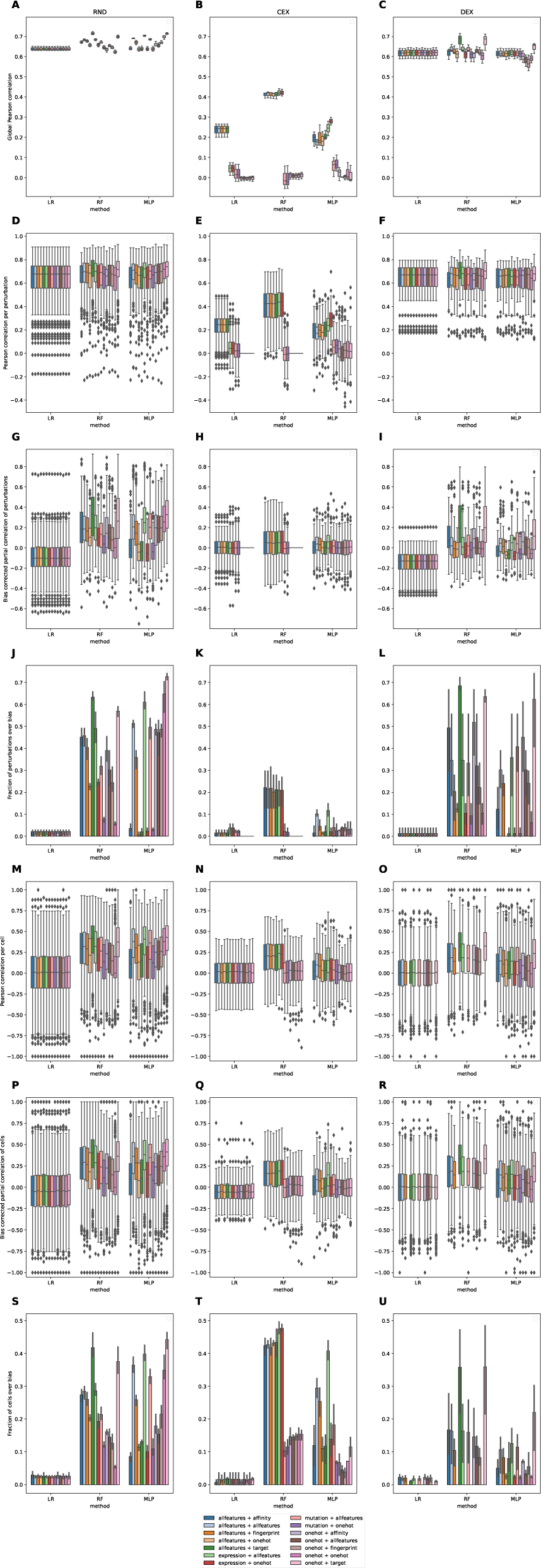
Model performance of predicting z-score drug sensitivity. Used evaluation metrics are labeled on the y-axis, used ML model is labeled on the x-axis, results are grouped based on splits (RND, CEX, GEX, and AEX, columns). Used cell line and perturbation feature pairs are color-coded (legend).

**Supplementary Table 1:** Target pathway enrichment for predictable drug sensitivities. Top 5 enriched pathways using target–correlation vectors as input for enrichment analysis. Correlations (raw or partial, Metrics column) were calculated between predicted and observed drug response (for CEX and DEX split, Split column), and aggregated drug-target wise. Reactome pathway enrichment was used with viper algorithm from decoupleR. The top 5 pathways are shown for each setup.

| Split | Metric | Rank | Reactome pathway |
| --- | --- | --- | --- |
| CEX | correlation | 1 | PROTEIN FOLDING |
| CEX | correlation | 2 | ASSOCIATION OF TRIC CCT WITH TARGET PROTEINS DURING BIOSYNTHESIS |
| CEX | correlation | 3 | TP53 REGULATES TRANSCRIPTION OF GENES INVOLVED IN CYTOCHROME C RELEASE |
| CEX | correlation | 4 | TP53 REGULATES TRANSCRIPTION OF CASPASE ACTIVATORS AND CASPASES |
| CEX | correlation | 5 | TP53 REGULATES TRANSCRIPTION OF CELL DEATH GENES |
| CEX | partial correlation | 1 | UPTAKE AND FUNCTION OF ANTHRAX TOXINS |
| CEX | partial correlation | 2 | UPTAKE AND ACTIONS OF BACTERIAL TOXINS |
| CEX | partial correlation | 3 | RAF ACTIVATION |
| CEX | partial correlation | 4 | SIGNALING BY MODERATE KINASE ACTIVITY BRAF MUTANTS |
| CEX | partial correlation | 5 | SIGNALING BY BRAF AND RAF FUSIONS |
| DEX | correlation | 1 | G1 PHASE |
| DEX | correlation | 2 | ONCOGENE INDUCED SENESCENCE |
| DEX | correlation | 3 | SENESCENCE ASSOCIATED SECRETORY PHENOTYPE SASP |
| DEX | correlation | 4 | OXIDATIVE STRESS INDUCED SENESCENCE |
| DEX | correlation | 5 | CYCLIN A:CDK2 ASSOCIATED EVENTS AT S PHASE ENTRY |
| DEX | partial correlation | 1 | SIGNALING BY NON RECEPTOR TYROSINE KINASES |
| DEX | partial correlation | 2 | DOWNREGULATION OF ERBB2:ERBB3 SIGNALING |
| DEX | partial correlation | 3 | GRB7 EVENTS IN ERBB2 SIGNALING |
| DEX | partial correlation | 4 | ERBB2 ACTIVATES PTK6 SIGNALING |
| DEX | partial correlation | 5 | SHC1 EVENTS IN ERBB2 SIGNALING |

## Footnotes

1 This is more just an example than an actual well-mapping downstream application since there is always some cell line and mouse data available on the drug of interest when starting clinical trials. However, a human patient is such a different environment, that it may not be a big stretch to say that it’s similar to a no-drug-data-available scenario.

