## Supplementary figures and images for "The EFFECT benchmark suite: measuring cancer sensitivity prediction performance - without the bias"

### Supplementary Figure 1

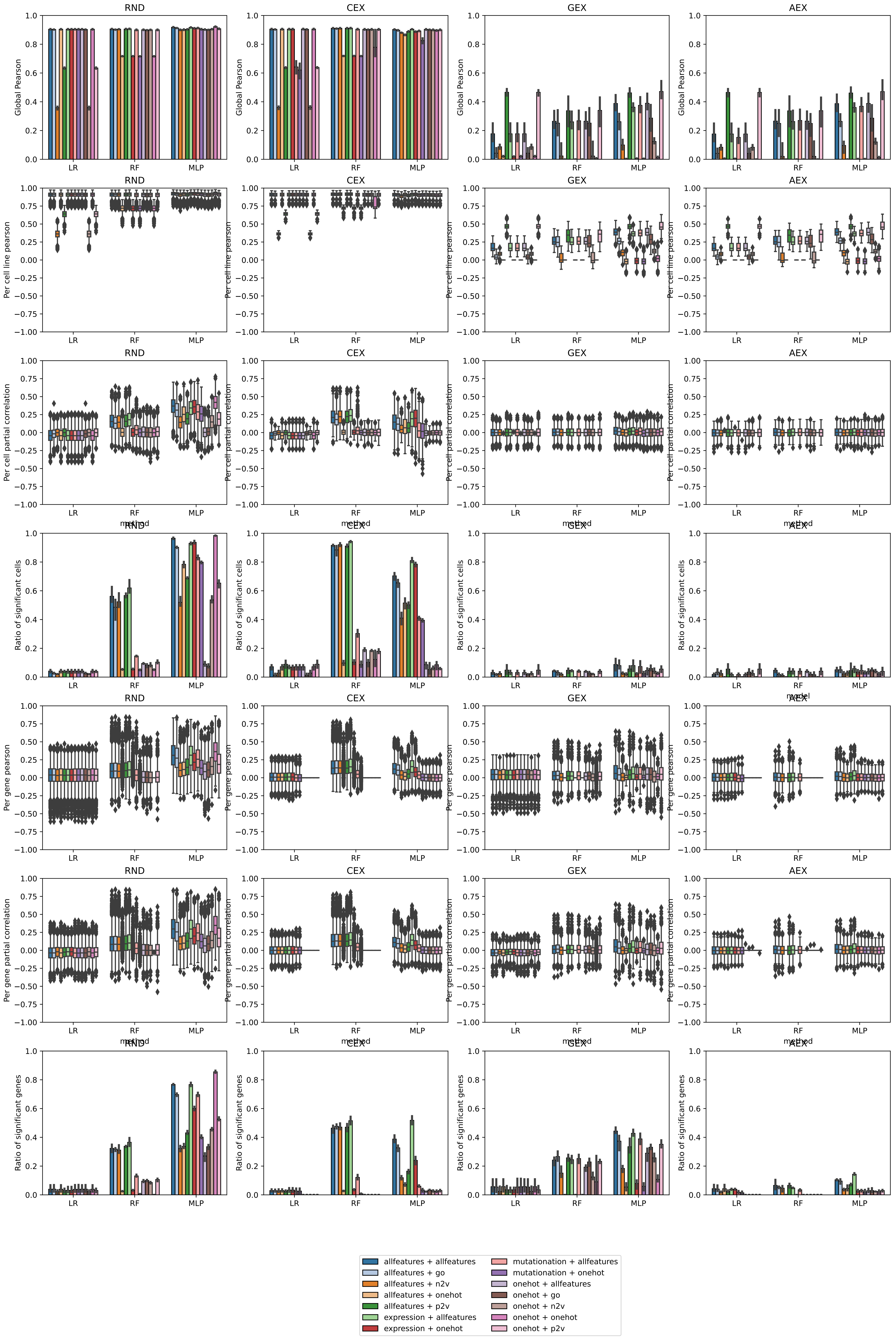

### Supplementary Figure 2

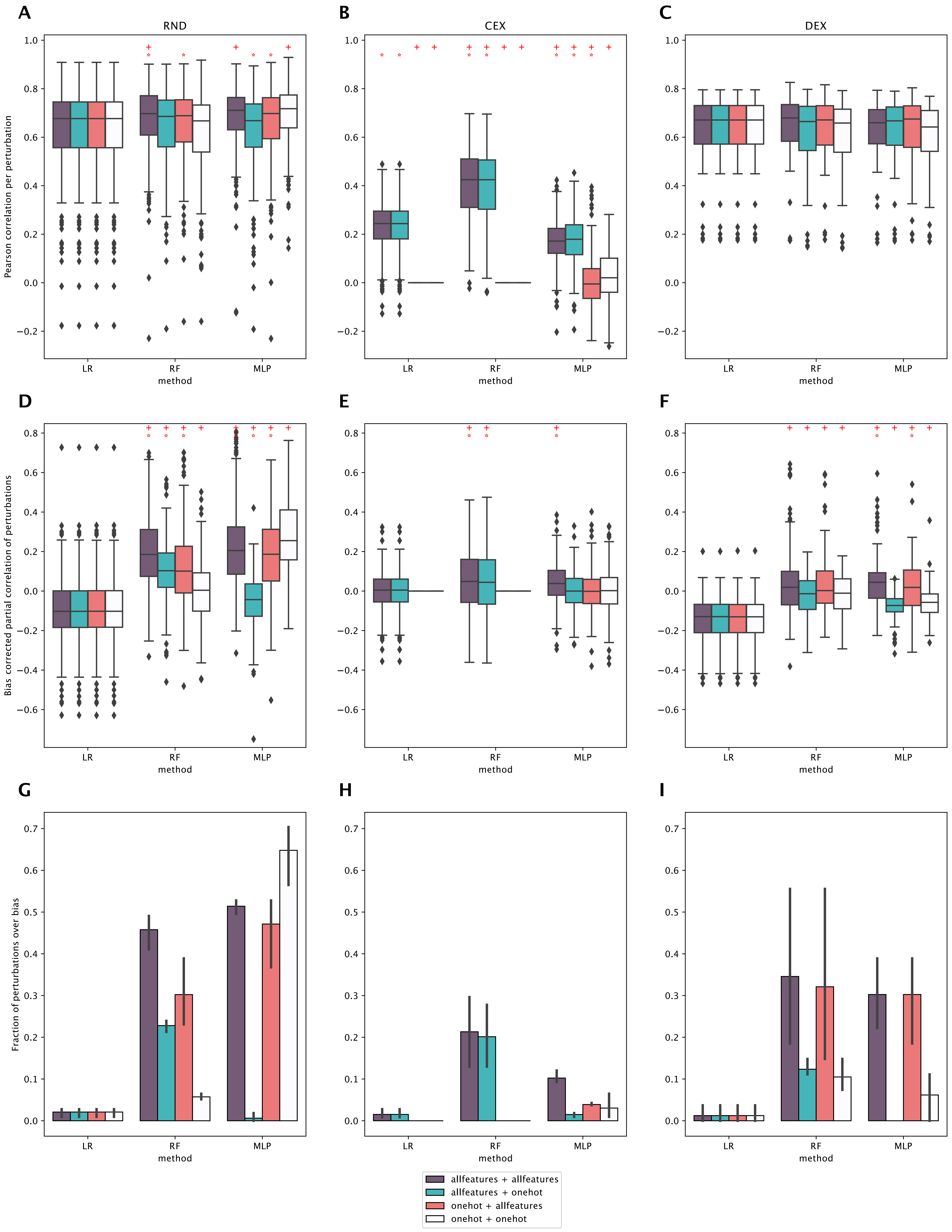

### Supplementary Figure 3

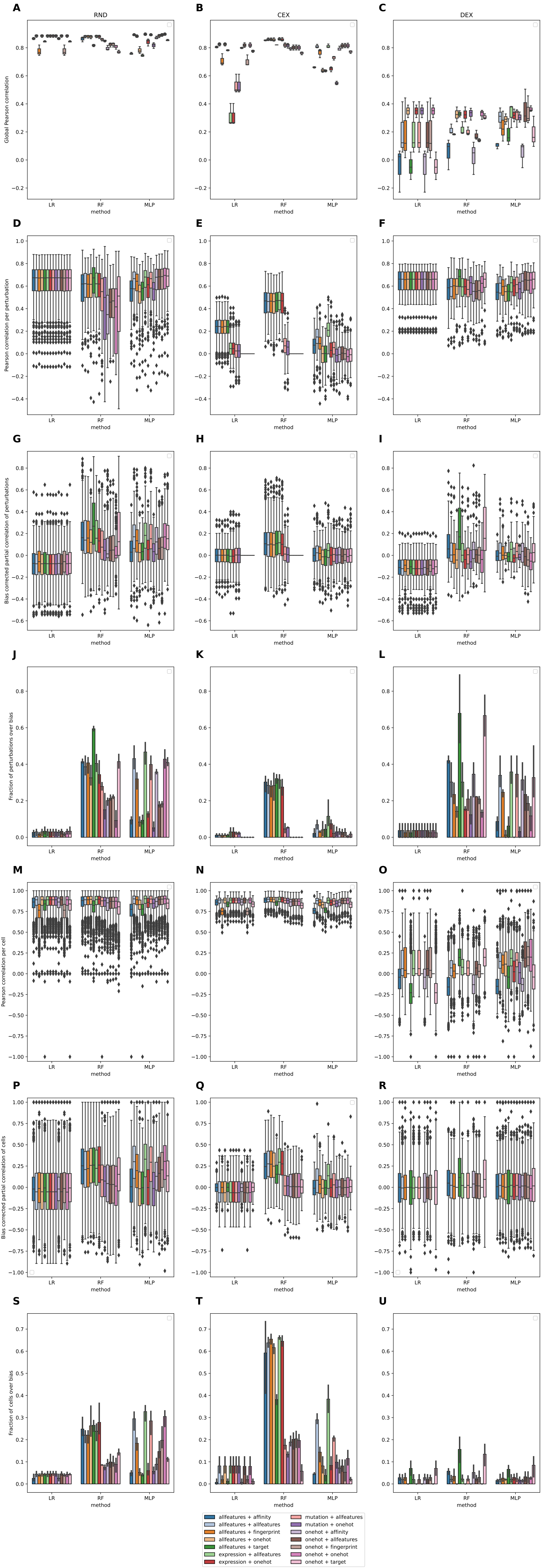

### Supplementary Figure 4

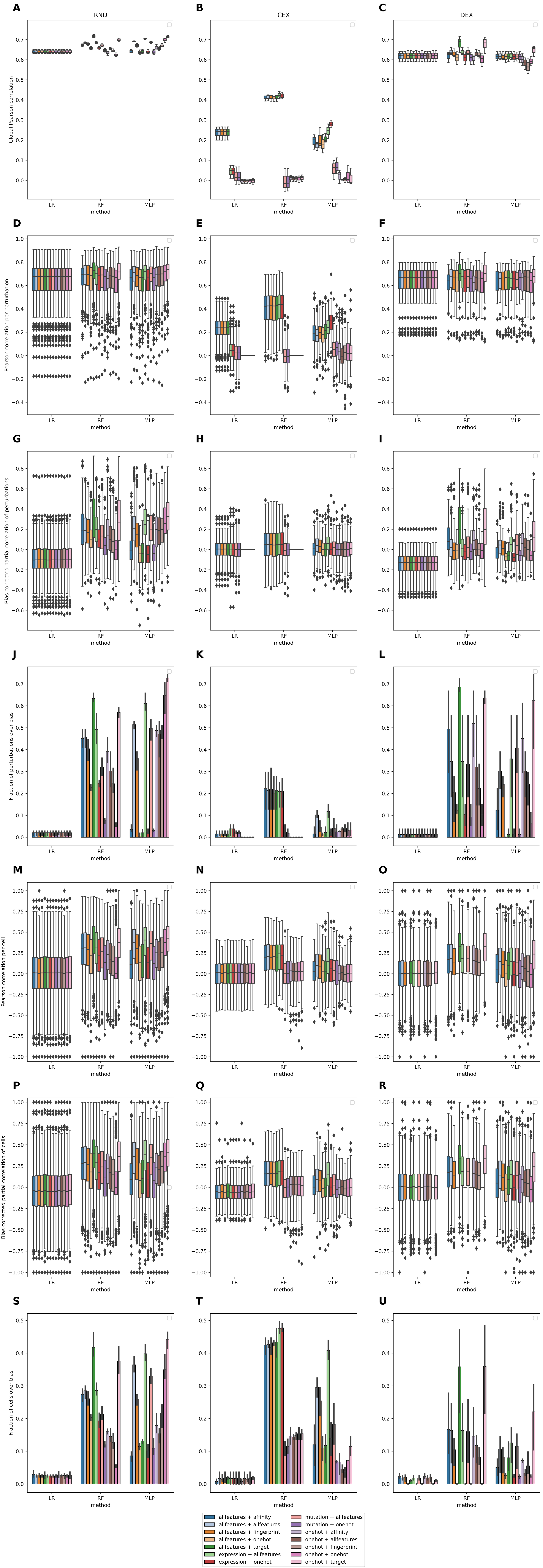
